# A modelling exercise to show why population models should incorporate distinct life histories of dispersers

**DOI:** 10.1101/402198

**Authors:** Jacques A. Deere, Ilona van den Berg, Gregory Roth, Isabel M. Smallegange

## Abstract

Dispersal is an important form of movement influencing population dynamics, species distribution, and gene flow between populations. In population models, dispersal is often included in a simplified manner by removing a random proportion of the population. Many ecologists now argue that models should be formulated at the level of individuals instead of the population-level. To fully understand the effects of dispersal on natural systems, it is therefore necessary to incorporate individual-level differences in dispersal behaviour in population models. Here we parameterised an integral projection model (IPM), which allows for studying how individual life histories determine population-level processes, using bulb mites, *Rhizoglyphus robini*, to assess to what extent dispersal expression (frequency of individuals in the dispersal stage) and dispersal probability affect the proportion of dispersers and natal population growth rate. We find that allowing for life-history differences between resident phenotypes and disperser phenotypes shows that multiple combinations of dispersal probability and dispersal expression can produce the same proportion of leaving individuals. Additionally, a given proportion of dispersing individuals results in different natal population growth rates. The results highlight that dispersal life histories, and the frequency with which disperser phenotypes occur in the natal population, significantly affect population-level processes. Thus, biological realism of dispersal population models can be increased by incorporating the typically observed life history differences between resident phenotypes and disperser phenotypes, and we here present a methodology to do so.

## Introduction

The movement of individuals is one of the key mechanisms shaping biodiversity (Travis and Dytham 1999; Jeltsch et al. 2013). An important form of movement is dispersal, which is any movement of individuals or propagules with potential for gene flow across space (Ronce 2007; Bonte et al. 2012). Dispersal influences the dynamics and persistence of populations, the distribution and abundance of species, the community structure, and the level of gene flow between populations (Brown and Kodric-Brown 1977; Dieckmann et al. 1999; Hanski 1999; Bowler and Benton 2005). In so doing, dispersal can fuel evolutionary processes such as local adaptation and speciation (Dieckmann et al. 1999). Currently, understanding dispersal behaviour is important to be able to predict how populations will respond to some of the most important threats to biodiversity, such as climate change, habitat loss and fragmentation, and the invasion of alien species (Bowler and Benton 2005; Clobert et al. 2009).

In the last decades, the drivers of dispersal have been the centre of many theoretical studies (Hamilton and May 1977; Johnson and Gaines 1990; Hanski 1999; Clobert et al. 2004). However, the dispersal process itself has gained less attention. Due to practical problems associated with the study of dispersal in the field, much of dispersal research has taken a theoretical approach (Bélichon et al. 1996; Bowler and Benton 2005). In population models, dispersal is often considered as a population-level process, in which a proportion of the population leaves (Clobert et al. 2004; Bowler and Benton 2005). In such models, individual-level differences are often ignored, which means that any individual within the population has the same probability of dispersing successfully. However, recently it has been argued that population ecology should shift its focus from formulating models at the population level to the level of individual organisms (Clark et al. 2011; Topping et al. 2015; Soudijn and Roos 2017); an argument echoed in the field of dispersal (Bonte et al. 2012). Although ecological patterns can be observed at the population level, it is the behaviour and demographic changes of individuals that shape the dynamics of the population (Clark et al. 2011; Soudijn and Roos 2017). So, in order to fully understand the effects of dispersal on natural systems, it is necessary to incorporate individual-level differences in dispersal behaviour in population models.

Empirical studies have shown that dispersing individuals typically have different demographic properties compared with residents (individuals that remain in the population) (Harrison 1980; Bélichon et al. 1996; O’Riain et al. 1996; Zera and Denno 1997; Deere et al. 2015). Dispersers and residents can differ in their morphology, physiology or in their behaviour (Bélichon et al. 1996; Clobert et al. 2009; Bonte et al. 2012). In many species, an individual property, e.g. locomotory movement, which potentially serves many important functions, is enhanced for dispersal (Phillips et al. 2006; Bonte et al. 2012). In other species, dispersers develop a morphology primarily as an adaptation for dispersal, e.g. the development of wings or structures which facilitate attachment to mobile vectors (Harrison 1980; Zera and Denno 1997; Diaz et al. 2000; Bonte et al. 2012). Species with a distinct disperser morphology are ideal study systems to investigate the individual-level effects and adaptive significance of dispersal in natural populations. For many of these species, each genotype has the potential to develop into a disperser or a resident (Harrison 1980; Clobert et al. 2004). The investment in a disperser morphology is costly if it requires resource investment, likely at the expense of investment into body condition or fecundity (Lemel et al. 1997; Bonte et al. 2012; Deere et al. 2015). Such investment costs, defined as “pre-departure costs”, arise during development prior to the actual dispersal event, although they may be deferred to later in the life-history as well (Bonte et al. 2012). However, not all individuals that invest in dispersal morphology leave their natal population, e.g. due to a behaviour or physiological component affecting dispersal propensity (Roff and Fairbairn, 2001), reduced patch accessibility (Clobert et al. 2001) or environmental conditions (Johnson 1969). In order to make the dispersal process in population models more realistic and its results more biologically relevant, an individual-based method that takes distinct dispersal life-histories and (un)successful dispersal into account is necessary. Here, we do so using an approach that incorporates these distinct dispersal life-histories into a population model structured by life stage. As a first step towards such an approach we focus on the natal population, where the dispersal process is initiated, and how dispersing individuals impact its dynamics. We apply the model to the invasive bulb mite (*Rhizoglyphus robini*, Claparède), the bulb mite is an ideal model system to study dispersal as it has a distinct dispersal stage within its life-history.

We model the system using Integral Projection Models (IPMs) as they comprise individual-level functions that describe demographic rates (thereby tracking fluctuations in population size and structure), they can be applied to species with complex demography, require relatively straightforward mathematical techniques from matrix calculus, and are closely and easily linked to field and experimental data (Easterling et al. 2000; Ellner and Rees 2006). To test our approach, we parameterise an IPM that incorporates a distinct dispersal life-history, where a dispersal stage is included in the life-cycle, and that incorporates an emigration component (where dispersers are able to leave the population). Within the IPM, each individual has the potential to develop into a *disperser phenotype* (i.e. develop into the dispersal stage) or a *resident phenotype* (i.e. do not develop into a dispersal stage), and each disperser phenotype has the option to leave the natal population (*disperser*) or remain in the natal population (*unsuccessful disperser*) (Fig. 1). The natal population will thus consist of resident phenotypes and unsuccessful dispersers. In this model, the individual-level costs of dispersal are then expressed by unsuccessful dispersers. It is important to note that unsuccessful dispersers in the context of this study refers to disperser phenotypes that do not emigrate from the population; not to disperser individuals that do not survive the transfer phase or fail to establish in a new habitat. To address our aim of understanding how distinct dispersal life histories play a role in natal population-level processes during a dispersal event, requires knowing not only the proportion of resident phenotypes and disperser phenotypes in the natal population, but also the proportion of disperser phenotypes that emigrate (dispersers). We will thus assess (i) the effects of the expression of a distinct disperser phenotype (frequency of individuals developing into disperser phenotypes) as well as the dispersal probability (probability that disperser phenotypes emigrate) on the proportion of dispersers (i.e. the disperser phenotypes that emigrate), which will allow us to tease apart the relative effects of the latter two dispersal components, and (ii) assess the knock-on consequences of disperser phenotype expression and actual dispersal on the growth rate of the natal population (Fig. 1).

**Figure 1.** Schematic indicating how a population-level approach and an individual-level approach differ in terms of the individuals that emigrate from a population. The blue blocks indicate individuals that stay in the natal population whilst grey blocks indicate individuals that leave the population.

## Methods

### Study system and life-history data

Bulb mites consist of five life stages, but develop an additional, sixth dispersal stage during development (called the deutonymph stage) under unfavourable environmental conditions (e.g. low temperature, humidity, food quality) (Diaz et al. 2000) (Fig. 2). The (juvenile) dispersal stage is non-feeding and occurs in both sexes. We use the same life-history data on female bulb mites as used in Deere et al. (2017) to parameterise the IPM; these data provide information on individuals that do and do not develop into the deutonymph stage as life histories differ between individuals that went through the deutonymph stage and those that did not (for detailed data collection see Deere et al. (2015). Data can be found in the figshare repository <u>10.6084/m9.figshare.1312875</u>).

**Figure 2.** Life cycle of the bulb mite in in the Disperser Phenotype Model (DPM), indicating the life stages and the vital rates. From the life cycle we calculated the survival (S) and fecundity (*F*) rates and the probability of growing and transitioning into the next stage (*G*). Dispersal consists of development into a dispersal phenotype (deutonymph) (β) and dispersal out of the population (*δ*); β describes the transition probability of developing from a protonymph to a deutonymph, *δ* describes the probability of a deutonymph leaving the population.

### Disperser Phenotype Model (DPM)

Our model, henceforth DPM, has a distinct dispersal stage, the deutonymph stage, which allows for emigration. Dispersal is a two-step process that consists of development into a disperser (deutonymph) and dispersal out of the population (Fig. 2). Deutonymph frequency, β, describes the transition probability of developing from a protonymph to a deutonymph. Dispersal probability, *δ*, describes the probability of a deutonymph leaving the population (Fig. 2). Together, they determine the proportion of individuals dispersing from a population. Whereas most models of dispersal assume *δ* = 1, our framework allows for the exploration of how variation in both affect populations.

We use the size- and stage-structured IPM developed by Deere et al. (2017) and we include an emigration component to the model. Briefly, an IPM tracks fluctuations in population size and structure based on individual-level processes. The IPM projects a new stage-size joint distribution based on the stage-size joint distribution in the previous time step (Easterling et al. 2000; Coulson 2012; Deere et al. 2017). For the DPM, in each time step individuals may survive, disperse, grow and produce new individuals. These processes are captured in a kernel, *K*, that projects joint distribution n_t_(*z,s*) of body size *z*, and stage *s*, at time *t*, to the new joint distribution at time *t*+1:

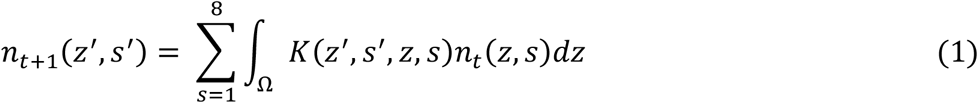

where Ω designates the range of individual sizes. The kernel, K, is composed of two parts,

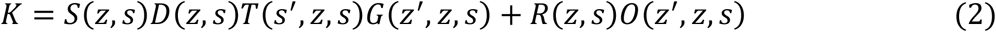

The first part describes how individuals grow, survive and move: S(*z,s*) is the survival probability to the next time step of an individual in size *z* and stage *s*; D(*z,s*) is the probability that an individual of size *z* in stage *s* stays in the population in the next time step; T(*s’,z,s*) is the probability that an individual in size *z* and stage *s* develops into the new stage *s’* and G(*z’,z,s*) is the probability that an individual in size *z* and stage *s* growths to size *z’*. The second part describes adult reproduction: R(*z,s*) is the number of offspring produced by an adult individual of size *z* in stage *s* and O(*z’,z,s*) is the probability that offspring produced by adults of size *z* are of size *z’* (i.e. the parent-offspring association). DPM equations (Table A1 equations 3.1 to 3.12), and parameter values used, can be found in the appendix.

The order of the functions in the kernel K represents the sequence of events in each time step, and the point at which the population is censused. This order is crucial for the outcome of the IPM (Rees et al. 2014). The order of events in the DPM is first survival, then dispersal, then transition of the individuals that do not disperse, followed by their growth. Dispersal is applied prior to transition in order to track all individuals that develop into the deutonymph stage and emigrate (the dispersers). Since dispersal is only possible for individuals in the deutonymph stage (i.e. dispersal stage), the probability to stay in the population is equal to 1 for all individuals in all stages other than the deutonymph stage. For all individuals in the deutonymph stage, the probability to stay in the population is 1 – *δ*.

### Models parameterization and analysis

The DPM was parameterised from life-history data of female bulb mites (see *Study system and life-history data*). Once parameterised, each function is then discretised into a matrix form by dividing the full size domain into very small-width discrete bins (‘mesh points’; see appendix for details). These discretised matrices describe the predicted transition rates and are used to build a projection matrix that approximates the DPM, so standard methods for analysing matrix population models can be used (Caswell 2001; Coulson 2012). Details on how the functions were parameterized can be found in the appendix. All analyses and simulations were performed in R version 3.0.2 (R Development Core Team 2013).

### Combination of disperser expression and dispersal probability

For the DPM we wanted to assess how an increase in deutonymph frequency (β) (i.e. (disperser phenotype expression) and dispersal probability (*δ*) would affect population-level processes. Within the DPM, β and *δ* were increased from 0-1 (at 0.01 increments) and for every combination of β and *δ* the proportion of individuals that disperse and population growth rate, λ_0_, were calculated. The proportion of individuals that disperse was calculated by integrating the stable stage distribution over the range of all sizes in the deutonymph stage and multiplying it with the dispersal probability *δ*. The stable stage distribution was calculated as the dominant right eigenvector of the projection matrix associated with each DPM (Easterling et al. 2000), λ_0_ was calculated as the dominant eigenvalue of the projection matrix associated to each DPM (Easterling et al. 2000).

## Results

When varying β and *δ* in the DPM there is a non-linear, joint effect of dispersal phenotype expression and dispersal probability on the proportion of dispersing individuals (Fig. 3) and population growth rate (λ_0_) (Fig. 4). If both β and *δ* increase then, in general, the proportion of dispersing individuals increases and λ_0_ decreases. Reciprocally, if both β and *δ* decrease then, in general, the proportion of dispersing individuals decreases and λ_0_ increases. When the changes in β and *δ* are opposed, any scenario (increase/decrease) in the proportion of dispersing individuals and λ_0_ can occur (Table 1).

**Table 1.** Response of population parameters to changes in dispersal parameters. Dots represent population parameter outcomes given various dispersal parameter scenarios. Black dots indicate expected outcomes, grey dots indicate unexpected outcomes (see main text for details).

| Dispersal parameters | Population parameters |  |  |  |
| --- | --- | --- | --- | --- |
| | PD↑ $\lambda_0$↓ | PD↓ $\lambda_0$↑ | PD↓ $\lambda_0$↓ | PD↑ $\lambda_0$↑ |
| $\beta$↑, $\delta$↑ | • | | | |
| $\beta$↓, $\delta$↓ | | • | | |
| $\beta$↑, $\delta$↓ | • | • | • | |
| $\beta$↓, $\delta$↑ | • | • | | • |

$\beta$ , disperser expression; $\delta$ , dispersal probability; PD, proportion of dispersers; $\lambda_0$ , population growth rate

**Figure 3.** Joint effect of increasing deutonymph frequency (β) and increasing dispersal probability (*δ*) on the proportion of dispersers. Dispersal probability and deutonymph frequency increase from 0 to 1 at 0.01 increments. Side bar indicates proportion of dispersers with proportion increasing from dark to light green. Coloured contour lines highlight a fixed proportion of dispersers, each proportion can be attained by a number of different *δ* and β values: 0.04 (yellow line), 0.08 (red line), 0.12 (blue line).

**Figure 4.** Joint effect of increasing deutonymph frequency (β) and increasing dispersal probability (*δ*) on population growth rate (λ_0_). Dispersal probability and deutonymph frequency increase from 0 to 1 at 0.01 increments. Side bar indicates λ_0_, with λ_0_ increasing from black to white. Coloured contour lines highlight a fixed proportion of dispersers (0.04, yellow line; 0.08, red line; 0.12, blue line), each proportion can result in a number of different λ_0_ values.

The joint effect of β and *δ* on the proportion of dispersers does not predict the joint effect of β and *δ* on λ_0_. Indeed, the contour lines of the proportion of dispersers intersect the contour lines of the λ_0_ (Fig. 3 & 4). As a consequence, for a fixed proportion of dispersing individuals, multiple λ_0_ values can occur (proportion of dispersing individuals: 0.04 (yellow line), 0.08 (red line), 0.12 (blue line); Fig. 3 & 4). Furthermore, with increasing proportions of dispersing individuals (yellow line < red line < blue line), the values of λ_0_ encompassed by each proportion of dispersing individuals, decreases (Fig. 4). Another consequence of the joint effect of β and *δ* is unexpected scenarios in the proportion of dispersing individuals and λ_0_ (grey dots in Table 1). For example, if β increases from 0.4 to 0.6 and *δ* decreases from 0.4 to 0.1, then the proportion of dispersing individuals decreases from 0.039 to 0.019. If β increases from 0.3 to 0.6 and *δ* decreases from 0.5 to 0.2, then λ_0_ decreases from 1.19 to 1.16 (Fig. 3 & 4). Vice versa, if β decreases and *δ* increases (in some specific proportion), then both the proportion of dispersing individuals and λ_0_ can increase. For example, the proportion of dispersing individuals increases from 0.045 to 0.066 when β decreases from 0.8 to 0.6, and *δ* increases from 0.2 to 0.5. In the case of λ_0_ there is an increase from 1.15 to 1.17 when β decreases from 0.7 to 0.2, and *δ* increases from 0.2 to 0.5 (Fig. 3 and 4).

The joint increase of β and *δ* can lead the population to extinction (i.e. λ_0_<1). Indeed, although the population growth rate is larger than 1 for almost all the parameter values, it is slightly smaller than 1 (λ_0_ = 0.99) when both the deutonymph frequency and the dispersal probability are equal to 1 (*δ* = β = 1). This high value of the population growth rate, even though all new-born individuals leave the population before reproducing, is a consequence of the high adult survival rate which maintains the population for a long time before it eventually goes extinct. When the adult survival rate is reduced, the population growth rate becomes smaller than 0.99 for larger values of β and *δ* (see Fig. S1).

## Discussion

The need to incorporate complexity and realism in modelling dispersal (Bonte et al. 2012; Travis et al. 2012; Travis et al. 2013) has prompted a move to incorporate more individual heterogeneity and individual-level costs when modelling dispersal (Bonte et al. 2012). Here we incorporated individual heterogeneity, in terms of distinct life-histories for disperser phenotypes and resident phenotypes, as well as the probability of disperser phenotypes to emigrate using an individual-based method. We applied our method to a system that has a distinct dispersal stage and assessed the effects of these distinct life-histories and emigration probability on population-level processes.

When modelling dispersal, dispersal rates are often applied to the whole population and the potential effect of dispersal to the natal population is in terms of the proportion of the individuals that leave. Given that the proportion of individuals that leave the population during a dispersal event is partly dependent on the number of individuals capable of dispersing within the population, we wanted to investigate the effect on the population of jointly manipulating the proportion of the disperser phenotypes in the population (β) and the dispersal probability (*δ*). Although we found a general trend for proportion of dispersers to increase (or decrease) and population growth rate to decrease (or increase) when both dispersal parameters increase (or decrease), when the two parameters increase or decrease in *opposite* directions, a number of different outcomes of the population parameters can occur (Table 1). The outcomes include scenarios which one would largely expect; an increase in the proportion of dispersers and a decrease in population growth (or vice versa). For example, when the proportion of dispersers increase, population growth rate will decrease as these individuals leave the population and so do not contribute to the population growth. However, there are also outcomes that we did not expect; scenarios where both the proportion of dispersers and population growth rate decrease or increase together. The scenario where the proportion of dispersers and population growth rate decrease occurs when β increases and *δ* decreases. An increase in the expression of deutonymphs results in more disperser phenotypes within the population, however, with lower dispersal probabilities, this would mean a reduced proportion of dispersers that emigrate. In the context of our system, disperser phenotypes are reliant on other insect species to disperse with disperser phenotypes attaching to their host via have a sucker plate on the dorsal part of their body (Diaz et al. 2000). As such, fewer or no insect species present, for example under colder or windier environment conditions across seasons, or even within the same day, will result in a reduced proportion of dispersers that emigrate. With fewer disperser phenotypes leaving the population, the individual-level costs of investing in dispersal (i.e. reduced size at maturity, reduced lifetime egg production, increased development time; Deere et al. 2015) are then borne out at the population level, thereby reducing population growth rate (Deere et al. 2017). The scenario where both the proportion of dispersers and population growth rate increase, occurs when β decreases and *δ* increases. Here, reducing β reduces the expression of deutonymphs within the population, whereas increasing *δ* results in a larger proportion of the disperser phenotypes leaving the population (i.e. an increase in the proportion of dispersers). Population growth rate will increase as fewer disperser phenotypes, with their associated individual-level costs of investing in dispersal, are left in the population and have a reduced effect on population growth rate. In all scenarios, expected and unexpected, the response is largely dependent on the ratio of the two dispersal parameters, highlighting that the population parameters are very responsive to the joint effect of the two dispersal parameters.

The biological implications of the joint effect of the two dispersal parameters are important. It shows how dispersal is more than just a probability of dispersing from a population: dispersal depends not only on the probability of phoresy (being able to disperse from the population) but also on the probability of developing into a disperser phenotype. What is more, the demographic consequences depend on both the probability of dispersing, as well as on the proportion of disperser phenotypes within the population that remain, as these disperser phenotypes carry a demographic cost of lower fecundity compared to those that have not invested in disperser morphology (residents) as a juvenile. The individual-level cost of investing in dispersal can be seen in the case where *δ* < 1 is compared to *δ* = 1, which is equivalent to comparing a scenario with and without individual-level differences respectively. For example, if we consider a population with a proportion of disperser phenotypes equal to 0.04 (yellow contour line, Fig 4), β = 0.86 and *δ* = 0.15 population growth rate equals 1.14. A model that does not account for individual-level differences but has an equal proportion of disperser phenotypes (i.e. *δ* = 1 and β = 0.86, yellow contour line), overestimates population growth rate (1.19). The demographic consequence of the proportion of disperser phenotypes in the population and the probability of dispersing is not restricted to species that invest in a distinct disperser phenotype. When individuals of a species disperse there is a cost to the individual (Bonte et al. 2012) and several environmental factors can impact individuals dispersing, such as reduced patch accessibility (Clobert et al. 2001) or environmental conditions (Johnson 1969; Cormont et al 2011; Kuussaari et al. 2016). For example, in the green-veined white butterfly (*Pieris napi*) increased flight ability, through investment in larger thoraces, is traded off against fecundity (Karlsson & Johansson 2008). However, given this investment in dispersal, there is no guarantee these individuals will disperse as butterfly dispersal is impacted by different weather variables. Increased temperatures increase butterfly dispersal propensity, but dispersal propensity decreases with increased cloud cover, rainfall and wind speed (Cormont et al 2011; Kuussaari et al. 2016). It follows that, to fully understand the influence of dispersing individuals on natal population processes, requires a detailed understanding of disperser life-histories and how the cost of investing in a disperser strategy is borne out at the population level, and to what extent disperser phenotypes are present in the population.

Another key finding is that different population structures (multiple combinations of β and *δ*) can give the same proportion of disperser individuals (Fig. 3); this is significant because population structure can have a large effect on population growth (Cameron et al. 2016; Smallegange et al. 2018). Previously, we have shown that, in the absence of emigration, distinct disperser phenotypes within the natal population have different life-histories to resident phenotypes (individuals unable to disperse) and so do not contribute to population processes in the same way (Deere et al. 2017). Additionally, juveniles and adults do not always respond the same way to changing environmental and population conditions and so differ in their contribution to populations dynamics (Coulson et al. 2001; Ozgul et al. 2012). Furthermore, the influence that the frequency of disperser phenotypes within the natal population and dispersal probability has on the proportion of dispersers can be seen as a natal-habitat induced effect. In our study the effects of the natal habitat on disperser phenotypes is clear. The expression of disperser phenotypes is dependent on the quality of the natal habitat with the expression of disperser phenotypes increasing as habitat quality decreases; however in other instances the effects may often be more subtle (e.g. maternal effects) (Benard and McCauley 2008). This effect on the proportion of dispersers can then further influence other processes such as genetic differentiation between populations (i.e. altering magnitude and/or symmetry of gene flow) (Benard and McCauley 2008). Indeed, Benard & McCauley (2008) highlight that environmentally induced asymmetries in the number or quality of dispersing individuals can lead to asymmetry in patterns of local adaptation. While our model is largely applicable to phoretic species, our findings are also relevant to actively dispersing species. In species showing active dispersal, dispersal is often dictated by a combination of dispersal ability (e.g. structural formation such as wings) and dispersal propensity (e.g. behaviour or physiological component), which would influence the proportion of dispersers. Moreover, it has been suggested that interactions between these traits can generate non-linear relationships between habitat condition and net dispersal rates (Benard and McCauley 2008). All in all, it is the combination of natal-habitat induced effects on disperser phenotypes and the population structure that will determine the proportion of dispersing individuals.

Local population dynamics and dispersal rates between populations determines the ecological dynamics of metapopulations. Indeed, dispersal rate has a large effect on metapopulation dynamics and has shown to be influential in the propensity for dispersal-induced stability and synchrony within metapopulation models (Abbott 2011) and patch-level asymmetry (Benard and McCauley 2008). Our results are limited to a natal population perspective, however this may ultimately effect metapopulation dynamics. A study by Altermatt and Ebert (2010) has shown that, when considering the origin and number of migrants, metapopulation functioning may differ to the patterns of the generally considered view of colonization-extinction dynamics of metapopulations (e.g. Hanski and Gaggiotti 2004). Altermatt and Ebert (2010) show that migrating stages occur in small and ephemeral habitat patches, contrary to colonization-extinction dynamics, and that these populations drive the metapopulation dynamics. In essence, they suggest that the focus should also be on where migrants colonizing a new habitat come from and in what numbers. Therefore, there is a need to accurately predict the proportion of dispersers that disperse from specific populations. We would further argue that the state of the natal population, given the presence of disperser phenotypes (e.g. population growth rate; see Deere et al. (2017)), would then also become important in the context of a metapopulation. Furthermore, models inform conservation management decisions. Many metapopulation models are used to make predictions that drive conservation management decisions and potentially highlight areas of data paucity (Hanski 1999; Clobert et al. 2001; Calabrese and Fagan 2004; Peterson and Freeman 2016; Morin et al. 2017). Modelling the dynamics of populations is by its very nature a simplified version of reality. However, as more complexity is added to models in a manner that improves the biological realism of how dispersal rates are implemented, then not only can model performance be improved in terms of their realism, but also in terms of the accuracy of their predictions, which ultimately is more informative when used in management decisions (Clobert et al. 2001; Peterson and Freeman 2016).

The importance of the joint effect of dispersal probability and the frequency of disperser phenotypes within the population that we show, indicates the potential importance of accounting for the frequency of disperser phenotypes within a population. We appreciate that our focus is on a specific study system, and do not suggest that our models are completely general in terms of their outcome. Rather we illustrate that the effects of dispersing individuals on natal populations are more than just a turnover of numbers. Given the importance of dispersal rates in (meta)populations, and that natural dispersal rates are being altered by human activities, how they are applied to populations then becomes vital. Only by identifying and including the individual-level costs disperser phenotypes in a population, and applying dispersal rates to those individuals, can the full extent of the effects of dispersal to natal populations, and potentially metapopulations, start to be realised.

## Acknowledgements

The work was funded by a MEERVOUD grant no. 836.13.001 and VIDI grant no. 864.13.005 from the Netherlands Organisation for Scientific Research to Isabel Smallegange.

## Author Contributions

JAD and IvdB conceptualized the study. JAD, IvdB and GR developed the models with input from IMS. JAD, IvdB, GR and IMS wrote and commented on the manuscript.

## Data Accessibility

Dataset used for model parametrisation can be found in the figshare repository 10.6084/m9.figshare.1312875.

## References

Abbott, K. C. 2011. A dispersal-induced paradox: synchrony and stability in stochastic metapopulations. Ecology Letters 14:1158–1169.

Altermatt, F., and D. Ebert. 2010. Populations in small, ephemeral habitat patches may drive dynamics in a Daphnia magna metapopulation. Ecology 91:2975–2982.

Bélichon, S., J. Clobert, and M. Massot. 1996. Are there differences in fitness components between philopatric and dispersing individuals? Acta Oecologia 17:503–517.

Benard, M. F., and S. J. McCauley. 2008. Integrating across Life-History Stages: Consequences of Natal Habitat Effects on Dispersal. The American Naturalist 171:553–567.

Bonte, D., H. Van Dyck, J. M. Bullock, A. Coulon, M. Delgado, M. Gibbs, V. Lehouck, et al. 2012. Costs of dispersal. Biological Reviews 87:290–312.

Bowler, D. E., and T. G. Benton. 2005. Causes and consequences of animal dispersal strategies: relating individual behaviour to spatial dynamics. Biological Reviews 80:205–225.

Brown, J. H., and A. Kodric-Brown. 1977. Turnover Rates in Insular Biogeography: Effect of Immigration on Extinction. Ecology 58:445–449.

Calabrese, J. M., and W. F. Fagan. 2004. A comparison-shopper’s guide to connectivity metrics. Frontiers in Ecology and the Environment 2:529–536.

Cameron, T. C., D. O’Sullivan, A. Reynolds, J. P. Hicks, S. B. Piertney, and T. G. Benton. 2016. Harvested populations are more variable only in more variable environments. Ecology and Evolution 6:4179–4191.

Caswell, H. 2001. Matrix population models: Construction, analysis, and interpretation (Second.). Sinauer Associates, Sunderland, Massachusetts.

Clark, J. S., D. M. Bell, M. H. Hersh, M. C. Kwit, E. Moran, C. Salk, A. Stine, et al. 2011. Individual-scale variation, species-scale differences: inference needed to understand diversity. Ecology Letters 14:1273–1287.

Clobert, J., E. Danchin, A. A. Dhondt, and J. D. Nichols, eds. 2001. Dispersal. Oxford University Press, New York.

Clobert, J., R. A. Ims, and F. Rousset. 2004. Causes, mechanisms and consequences of dispersal. Pages 307–335 in I. Hanski and O. E. Gaggiotti, eds. Ecology, Genetics and Evolution of Metapopulations. Elsevier Academic Press Inc, Amsterdam.

Clobert, J., J. F. Le Galliard, J. Cote, S. Meylan, and M. Massot. 2009. Informed dispersal, heterogeneity in animal dispersal syndromes and the dynamics of spatially structured populations. Ecology Letters 12:197–209.

Coulson, T. 2012. Integral projections models, their construction and use in posing hypotheses in ecology. Oikos 121:1337–1350.

Coulson, T., E. A. Catchpole, S. D. Albon, B. J. T. Morgan, J. M. Pemberton, T. H. Clutton-Brock, M. J. Crawley, et al. 2001. Age, Sex, Density, Winter Weather, and Population Crashes in Soay Sheep. Science, New Series 292:1528–1531.

Cormont, A., A. H. Malinowska, O. Kostenko, V. Radchuk, L. Hemerik, M. F. WallisDeVries, and J. Verboom. 2011. Effect of local weather on butterfly flight behaviour, movement, and colonization: significance for dispersal under climate change. Biodiversity and Conservation 20:483–503.

Deere, J. A., T. Coulson, S. Cubaynes, and I. M. Smallegange. 2017. Unsuccessful dispersal affects life history characteristics of natal populations: The role of dispersal related variation in vital rates. Ecological Modelling 366:37–47.

Deere, J. A., T. Coulson, and I. M. Smallegange. 2015. Life History Consequences of the Facultative Expression of a Dispersal Life Stage in the Phoretic Bulb Mite (Rhizoglyphus robini). PLoS ONE 10:e0136872.

Diaz, A., K. Okabe, C. J. Eckenrode, M. G. Villani, and B. M. Oconnor. 2000. Biology, ecology, and management of the bulb mites of the genus Rhizoglyphus (Acari?: Acaridae). Experimental & Applied Acarology 24:85–113.

Dieckmann, U., B. O’Hara, and W. Weisser. 1999. The evolutionary ecology of dispersal. Trends Ecol Evol 14.

Easterling, M. R., S. P. Ellner, and P. M. Dixon. 2000. Size-specific sensitivity: Applying a new structured population model. Ecology 81:694–708.

Ellner, S. P., and M. Rees. 2006. Integral Projection Models for Species with Complex Demography. The American Naturalist 167:410–428.

Hamilton, W. D., and R. M. May. 1977. Dispersal in stable habitats. Nature 269:578–581.

Hanski, I. 1999. Metapopulation ecology. Oxford University Press, New York.

Hanski, I., and O. E. Gaggiotti, eds. 2004. Ecology, Genetics and Evolution of Metapopulations. Elsevier Academic Press Inc, Amsterdam.

Harrison, R. G. 1980. Dispersal Polymorphisms in Insects. Annual Review of Ecology and Systematics 11:95–118.

Jeltsch, F., D. Bonte, G. Pe’er, B. Reineking, P. Leimgruber, N. Balkenhol, B. Schroder, et al. 2013. Integrating movement ecology with biodiversity research - exploring new avenues to address spatiotemporal biodiversity dynamics. Movement Ecology 1:6.

Johnson, C. G. 1969. Insect migration and dispersal by flight. Methuen, London.

Johnson, M. L., and M. S. Gaines. 1990. Evolution of Dispersal: Theoretical Models and Empirical Tests Using Birds and Mammals. Annual Review of Ecology and Systematics 21:449–480.

Karlsson, B., and A. Johansson. 2008. Seasonal polyphenism and developmental trade-offs between flight ability and egg laying in a pierid butterfly. Proceedings of the Royal Society B: Biological Sciences 275:2131–2136.

Kuussaari, M., S. Rytteri, R. K. Heikkinen, J. Heliölä, and P. von Bagh. 2016. Weather explains high annual variation in butterfly dispersal. Proceedings of the Royal Society B: Biological Sciences 283:20160413.

Lemel, J.-Y., S. Belichon, J. Clobert, and M. E. Hochberg. 1997. The evolution of dispersal in a two-patch system: some consequences of differences between migrants and residents. Evolutionary Ecology 11:613–629.

Morin, D. J., A. K. Fuller, A. Royle, and C. Sutherland. 2017. Model-based estimators of density and connectivity to inform conservation of spatially structured populations. Ecosphere 8:e01623.

O’Riain, M. J., J. U. M. Jarvis, and C. G. Faulkes. 1996. A dispersive morph in the naked mole-rat. Nature 380:619.

Ozgul, A., T. Coulson, A. Reynolds, T. C. Cameron, and T. G. Benton. 2012. Population Responses to Perturbations: The Importance of Trait-Based Analysis Illustrated through a Microcosm Experiment. American Naturalist 179:582–594.

Peterson, J. T., and M. C. Freeman. 2016. Integrating modelling, monitoring, and management to reduce critical uncertainties in water resource decision making. Journal of Environmental Management 183:361–370.

Phillips, B. L., G. P. Brown, J. K. Webb, and R. Shine. 2006. Invasion and the evolution of speed in toads. Nature 439.

R Core Team. 2013. R: A language and environment for statistical computing. R Foundation for Statistical Computing, Vienna, Austria. URL http://www.R-project.org/.

Rees, M., D. Z. Childs, and S. P. Ellner. 2014. Building integral projection models: a user’s guide. Journal of Animal Ecology 83:528–545.

Roff, D.A., Fairbairn, D.J., 2001. The genetic basis of dispersal and migration, and its consequences for the evolution of correlated traits. In: Clobert, J., Danchin, E., Dhondt, A.A., Nichols, J.D. (Eds.), Dispersal. Oxford University Press Inc., NewYork, pp. 191–202.

Ronce, O. 2007. How does it feel to be like a rolling stone? Ten questions about dispersal evolution. Annual Review of Ecology Evolution and Systematics 38:231–253.

Smallegange, I. M., R. E. Fernandes, and J. C. Croll. 2018. Population consequences of individual heterogeneity in life histories: overcompensation in response to harvesting of alternative reproductive tactics. Oikos 127:738–749.

Soudijn, F. H., and A. M. Roos. 2017. Approximation of a physiologically structured population model with seasonal reproduction by a stage-structured biomass model. Theoretical Ecology 10:73–90.

Topping, C. J., H. F. Alrøe, K. N. Farrell, and V. Grimm. 2015. Per Aspera ad Astra: Through Complex Population Modeling to Predictive Theory. The American Naturalist 186:669–674.

Travis, J. M. J., M. Delgado, G. Bocedi, M. Baguette, K. Barton, D. Bonte, I. Boulangeat, et al. 2013. Dispersal and species’ responses to climate change. Oikos 122:1532–1540.

Travis, J. M. J., and C. Dytham. 1999. Habitat persistence, habitat availability and the evolution of dispersal. Proceedings of the Royal Society of London B: Biological Sciences 266:723–728.

Travis, J. M. J., K. Mustin, K. a. Bartoń, T. G. Benton, J. Clobert, M. M. Delgado, C. Dytham, et al. 2012. Modelling dispersal: an eco-evolutionary framework incorporating emigration, movement, settlement behaviour and the multiple costs involved. Methods in Ecology and Evolution 3:628–641.

Zera, A. J., and R. F. Denno. 1997. Physiology and ecology of dispersal polymorphism in insects. Annual Review of Entomology 42:207–230.

