## appendix for "A modelling exercise to show why population models should incorporate distinct life histories of dispersers"

### Appendix A

This online appendix accompanies the paper "A modelling exercise to show why population models should incorporate distinct life-histories of dispersers" by Jacques A. Deere, Ilona van den Berg, Gregory Roth and Isabel M. Smallegange.

The first section describes the methods used to parameterize the character demography functions and calculation of the mesh points used in the DPM. We also give the parameter estimates for all the character-demography functions for the DPM. This is followed by the DPM equations used in the model which can be found in table A1. Figure S1 highlights how population growth rate, when varying β and $\delta$, is impacted when adult survival rate is altered.

#### *Parameter estimation*

Life-history data on female bulb mites taken from Deere et al. (2015) were used to parameterise the DPM. We estimated the parameters using statistical models for the five character-demography functions: the survival function (1), transition function (2), growth function (3), reproduction function (4) and the parent-offspring association function (5) (Fig. 2 in main text).

*Non-dispersers*

Parameters were estimated following the method used by Smallegange et al. (2014). Function parameters were estimated using the following statistical models (summarized in Table A1): (1) Survival - generalised linear mixed model (GLMMs) with binomial error structure; (2) Transition rates – GLMM with binomial error structure; (3) Growth – GLMM with a Gaussian error structure; (4) Fertility rate (Reproduction) – GLMM with a Gaussian error structure; (5) Parent-offspring association kernel – generalised linear model (GLM) with a Gaussian error structure. In all cases body length and body length squared were linear predictors and, with the exception of the parent-offspring association function where we fitted a GLM, mite identity was included as a random factor. The response variables for the five functions were: (1) Survival – from time *t* to time *t*+1 (this is binary and set as 0 or 1), (2) Transition - probability of growing to the next stage at time *t*+1, $\gamma_{s,t+1}$ (see below), (3) Growth - mean and variance in body size at time *t*+1, (4) Fertility - the number of eggs produced at time *t*+1, and (5) Parent-offspring association - mean and variance in size of eggs produced at time *t*+1 by each individual at time *t*. In the case of eggs their size at time *t*+1 equalled their size at time *t* as eggs do not increase in size.

During data collection it was not always possible to locate each individual every day. As such, the days where an individual was not seen but was still alive (i.e. observed alive the next day), body length was estimated (not including these observations would result in an underestimation of the survival function). The missing values were filled in by using the Gompertz function to estimate female body length at age *a* (Smallegange et al. 2014):

$$z_a = z_\infty e^{-e^{-k(a-a_0)}} \qquad \text{(Eq. S1)}$$

where $Z_a$ is body length (mm) at age *a* (days), $Z_\infty$ is the mean maximum length (mm, at $a = \infty$), *k* is the instantaneous growth rate at age $a_0$ , and $a_0$ is the inflection point of the curve and the age at which absolute growth rate begins to decline.

In the case of function (2), we used duration in the life stage as an indicator of the transitioning to the next stage (Smallegange et al. 2014); i.e. time spent in the current stage depends on the probability of growing to the next stage (Caswell 2001). Therefore, $\gamma_{s,t+1}$ is given by $\gamma_{s,t+1} = 1/d_{s,t}$, where $d_{s,t}$ is the number of days that an individual still has to spend in stage $s$. This means that $d_s$ equals the total duration of stage $s$ on the first day that a female is in stage $s$, and that when an individual develops from stage $s$ into stage $s$+1 at time $t$+1, $d_s$=1 so that $\gamma_{s,t+1}$ =1.

For function (3) and (5) the minimal model, which generated the predictors of mean size at time $t$+1, was utilised to generate the parameters for the variance around the mean size at $t$+1 by taking the squared residuals and fitting them against a statistical function of the same form as the mean size to estimate the variance in size at time $t$+1. The growth (3) and parent-offspring association functions (5) were then constructed using the equation following Easterling et al. (2000):

$$y_i = \frac{1}{\sqrt{2\pi\sigma_i}} e^{\frac{-(z' - \mu_i)^2}{2\sigma_i^2}} \qquad \text{(Eq. S2)}$$

where $y_i$ is either the growth or parent-offspring association function, $\mu_i$ describes the mean effect of the significant predictors on growth or parent-offspring association, and $\sigma_i$ describes the squared residuals around $\mu_i$.

For details on construction of the DPM from these parameters see main text. Note that mite identity was included in the statistical analyses (except in the parent-offspring association function) but was not modelled within the DPM.

*Dispersers*

In the case of dispersers the rates for eggs and larvae were the same as for the non-dispersers (see Deere et al. (2015) and Fig. 2 in main text). However, protonymphs can also develop into a deutonymph and we estimated the probability that a protonymph develops into a deutonymph, tritonymph or stays as a protonymph using a multinomial logisitic (which generated the three transition probabilities). For the multinomial logistic the linear predictor was body length at time *t*, the response variable was stage at time *t*+1 and the reference level was set as the protonymph stage. This gives the probability of developing into a tritonymph, a deutonymph or remaining as a protonymph as a function of individual size. As such, the regression coefficients $\beta\mu$ are the log of the ratio of the two probabilities of developing into a tritonymph or deutonymph over staying in the protonymph stage (the reference level/choice). For example, if $\beta\mu$ represents the effect of $\mu$ (size), we expect that for one unit change in $\mu$, the relative risk of developing into a tritonymph over staying a protonymph will increase by $\exp(\beta\mu)$. The multinomial logistic analyses were performed in R (version 3.0.2) using the 'mlogit' package (R Development Core Team 2013). All other parameter estimates were calculated in the same way, and using the same analyses, as those for non-dispersing individuals.

In all statistical analyses a model simplification procedure was used. The full model was fitted, after which the least significant term from the highest order interaction downwards was identified and removed if the removal resulted in an insignificant increase in deviance. The full and reduced models are shown in Table A1. Significance of simplified models was assessed by performing a likelihood ratio test. The likelihood ratio (Λ) is calculated as $\Lambda = 2(LL_f - LL_j)/(p_f - p_j)$; where $LL_i$ is the log-likelihood of the full model and $LL_j$ is the log-likelihood of the reduced model ($j$ = r) or constant-only model ($j = c$). $p_i$ is the number of estimable parameters in the full and $p_j$ is the number of estimable parameters in the reduced and constant-only model. The likelihood ratio is $X_{\nu}^2$ distributed, where $\nu$ is the difference in number of estimable parameters. The random factor was never removed during model simplification. Model assumptions and homoscedacity were confirmed by inspection of probability plots and error structures. All analyses were performed in R (version 3.0.2) with models fitted by maximum likelihood in the ‘lme4’ package (R Development Core Team 2013).

***Mesh point calculation***

Mesh points were created by dividing the size domain of each stage into very small-width discrete bins. A number of different bin sizes were used and results compared, this was done as an increase in the number of mesh points increases the numerical accuracy of the approximation (Ellner and Rees 2006). The body size domain of each stage was eventually divided into 50 size bins as a higher number of bins did not produce notably different results. Transition rates for the midpoint of two adjacent mesh points were estimated for each stage class. In the NM, the final matrix size was 250X250 (50 bins x 5 stages = 250 mesh points); whereas in the DPM the final matrix size was 400X400 (50 bins x 8 stages = 400 mesh points). The DPM takes into account the different number of life stages of dispersers and non-dispersers as well two sets of tritonymph and adult life stages (tritonymphs and adults without dispersal stage, and tritonymphs and adults with dispersal stage) into a single IPM, hence there are eight stages in the final matrix and not five (Fig. 2 in main text).

#### ***Parameter values of character-demography functions***

In the functions below, B denotes body length (mm) and sample size n is given in between brackets for each fitted function. E - egg; L – larva; P – protonymph; D – deutonymph; P/D – protonymph and deutonymph combined; T – tritonymph; TP – tritonymph from non-disperser; TD – tritonymph from disperser; A – adult; AP – Adult from non-disperser; AD – adult from disperser.

Survival rates for the DPM (fraction per day)

E: $y = 0.956$ ($n$ = 297); L: $y = 0.999$ ($n$ = 112); P: $y = 0.910$ ($n$ = 166); D: $y = 0.999$ ($n$ = 426); $T_P$: $y = 0.999$ ($n$ = 132); $T_D$: $y = \frac{1}{1+\frac{1}{e^{(-0.4175+6.9435B)}}}$ ($n$ = 119); $A_P$: $y = 0.999$ ($n$ = 115); $A_D$: $y = 0.933$ ($n$ = 60).

Life stage transition rates for the DPM (fraction per day)

E→L: $y = \frac{1}{1+\frac{1}{e^{(-1.437+8.674B)}}}$ ($n$ = 97); L→P: $y = \frac{1}{1+\frac{1}{e^{(-6.933 + 29.429B)}}}$ ($n$ = 47);

P→D: $y = \dfrac{1}{1 + \dfrac{1}{e^{(-2.601 + (-5.673)B)}}}$ ($n$ = 137); P→T: $y = \dfrac{1}{1 + \dfrac{1}{e^{(-11.220 + 26.235B)}}}$ ($n$ = 137); D→T: $y = \dfrac{1}{1 + \dfrac{1}{e^{(55.05 - 385.54B + 654.02B^2)}}}$ ($n$ = 155); $T_P$→$A_P$: $y = \dfrac{1}{1 + \dfrac{1}{e^{(-6.703 + 13.100B)}}}$ ($n$ = 76); $T_D$→$A_D$: $y = \dfrac{1}{1 + \dfrac{1}{e^{(-6.275 + 14.933B)}}}$ ($n$ = 45).

Reproduction rate for the DPM (no. per day)

$A_P$: $y = 0.5(-18.446 + 35.209B)$ ($n$ = 190); $A_D$: $y = 0.5(-13.592 + 33.892B)$ ($n$ = 172)

Mean growth rates for the DPM (when staying in the same life stage) (mm)

E: $y = L$ ($n$ = 65); L: $y = 0.11739 + 0.64316B$ ($n$ = 29); P: $y = 0.0772 + 0.904B$ ($n$ = 39); D: $y = L$ ($n$ = 153); $T_P$: $y = 0.0776 + 0.9538B$ ($n$ = 44); $T_D$: $y = -0.0772 + 1.3570B$ ($n$ = 23); $A_P$: $y = 0.3977 + 0.5359B$ ($n$ = 215); $A_D$: $y = 0.2816 + 0.6355B$ ($n$ = 238)

Variance in growth rates for the DPM (when staying in the same life stage) ($mm^2$)

E: $y = 0.0001$ ($n$ = 65); L: $y = -0.0008 + 0.0050B$ ($n$ = 29); P: $y = -0.0007 + 0.0040B$ ($n$ = 39); D: $y = 0.0001$ ($n$ = 153); $T_P$: $y = 0.0039 - 0.0042B$ ($n$ = 44); $T_D$: $y = -0.0044 - 0.0060B$ ($n$ = 23); $A_P$: $y = 0.0009 - 0.0004B$ ($n$ = 215); $A_D$: $y = 0.0014 - 0.0016B$ ($n$ = 238)

Parent-offspring association function (mean offspring-mother difference) for the DPM (mm)

$A_P$: $y = 0.1638$ ($n$ = 96); $A_D$: $y = 0.1689$ ($n$ = 175)

Variance around parent-offspring association function for the DPM ($mm^2$)

$A_P$: $y = 0.00008$ ($n$ = 96); $A_D$: $y = 0.0001$ ($n$ = 175)

**Table A1.** The DPM equations are constructed from the five statistical demography functions: Survival $\boldsymbol{S}(\boldsymbol{z},\boldsymbol{s},\boldsymbol{t})$, Transition $\boldsymbol{T}(\boldsymbol{s}'\,|\,\boldsymbol{z},\boldsymbol{s},\boldsymbol{t})$, Growth $\boldsymbol{G}(\boldsymbol{z}'\,|\,\boldsymbol{z},\boldsymbol{s}',\boldsymbol{t})$, Reproduction $\boldsymbol{R}(\boldsymbol{z},\boldsymbol{s},\boldsymbol{t})$ and Offspring inheritance $\boldsymbol{O}(\boldsymbol{z}'\,|\,\boldsymbol{z},\boldsymbol{s},\boldsymbol{t})$ and the dispersal matrix $\boldsymbol{D}(\boldsymbol{t})$. The equations calculate the number of females in each stage s at time t which is described by $\boldsymbol{n}(\boldsymbol{z},\boldsymbol{s},\boldsymbol{t})$ with the $\boldsymbol{R}(\boldsymbol{z},\boldsymbol{s},\boldsymbol{t})$ and the $\boldsymbol{O}(\boldsymbol{z}'\,|\,\boldsymbol{z},\boldsymbol{s},\boldsymbol{t})$ functions zero for all non-adult stages as only adults reproduce.

| | Life stage Equation | Description |
|---|---|---|
| (3.1) | $n(z',1,t+1) = \int O(z'\|z,5,t)R(z,5,t)n(z,5,t)dz + \int O(z'\|z,8,t)R(z,8,t)n(z,8,t)dz$ | Egg production by non-dispersal and dispersal adults |
| (3.2) | $\left.\begin{aligned} n(z',s+1,t+1) &= \int_{\Omega s} G(z'\|z,s+1,t)T(s+1\|z,s,t)S(z,s,t)n(z,s,t)dz \\ n(z',s,t+1) &= \int_{\Omega s} G(z'\|z,s,t)T(s\|z,s,t)S(z,s,t)n(z,s,t)dz \end{aligned}\right\} 1 \leq s \geq 2$ | Eggs and Larvae developing into the next stage and staying in the same stage |
| (3.3) | $n(z',3,t+1) = \int G(z'\|z,3,t)T(3\|z,3,t)S(z,3,t)n(z,3,t)dz$ | Non-dispersal Protonymphs staying Protonymphs |
| (3.4) | $n(z',6,t+1) = \int G(z'\|z,6,t)T(6\|z,3,t)S(z,6,t)n(z,3,t)dz$ | Deutonymphs developing from Protonymphs |
| (3.5) | $n(z',4,t+1) = \int G(z'\|z,4,t)T(4\|z,3,t)S(z,4,t)n(z,3,t)dz$ | Non-dispersal Tritonymphs developing from Protonymphs |
| (3.6) | $n(z',4,t+1) = \int G(z'\|z,4,t)T(4\|z,4,t)S(z,4,t)n(z,4,t)dz$ | Non-dispersal Tritonymphs staying Tritonymphs |

(Table 1 cont.)

| | Life stage Equation | Description |
|---|---|---|
| (3.7) | $n(z',6,t+1) = \int G(z'\|z,6,t)T(6\|z,6,t)D(6,t)S(z,6,t)n(z,6,t)dz$ | Deutonymphs staying Deutonymphs |
| (3.8) | $n(z',7,t+1) = \int G(z'\|z,7,t)T(7\|z,6,t)D(6,t)S(z,7,t)n(z,6,t)dz$ | Dispersal Tritonymphs developing from Deutonymphs |
| (3.9) | $n(z',8,t+1) = \int G(z'\|z,8,t)T(8\|z,7,t)S(z,8,t)n(z,7,t)dz$ | Dispersal adults developing from dispersal Tritonymphs |
| (3.10) | $n(z',7,t+1) = \int G(z'\|z,7,t)T(7\|z,7,t)S(z,7,t)n(z,7,t)dz$ | Dispersal Tritonymph staying Tritonymphs |
| (3.11) | $n(z',5,t+1) = \int G(z'\|z,5,t)T(5\|z,5-1,t)S(z,5-1,t)n(z,5-1,t)dz + \int G(z\|z,5,t)S(z,5,t)n(z,5,t)dz$ | Non-dispersal adults developing from non-dispersal Tritonymphs and surviving non-dispersal adults |
| (3.12) | $n(z',8,t+1) = \int G(z'\|z,8,t)T(8\|z,8-1,t)S(z,8-1,t)n(z,8-1,t)dz + \int G(z\|z,8,t)S(z,8,t)n(z,8,t)dz$ | Dispersal adults developing from dispersal Tritonymphs and surviving dispersal adults |

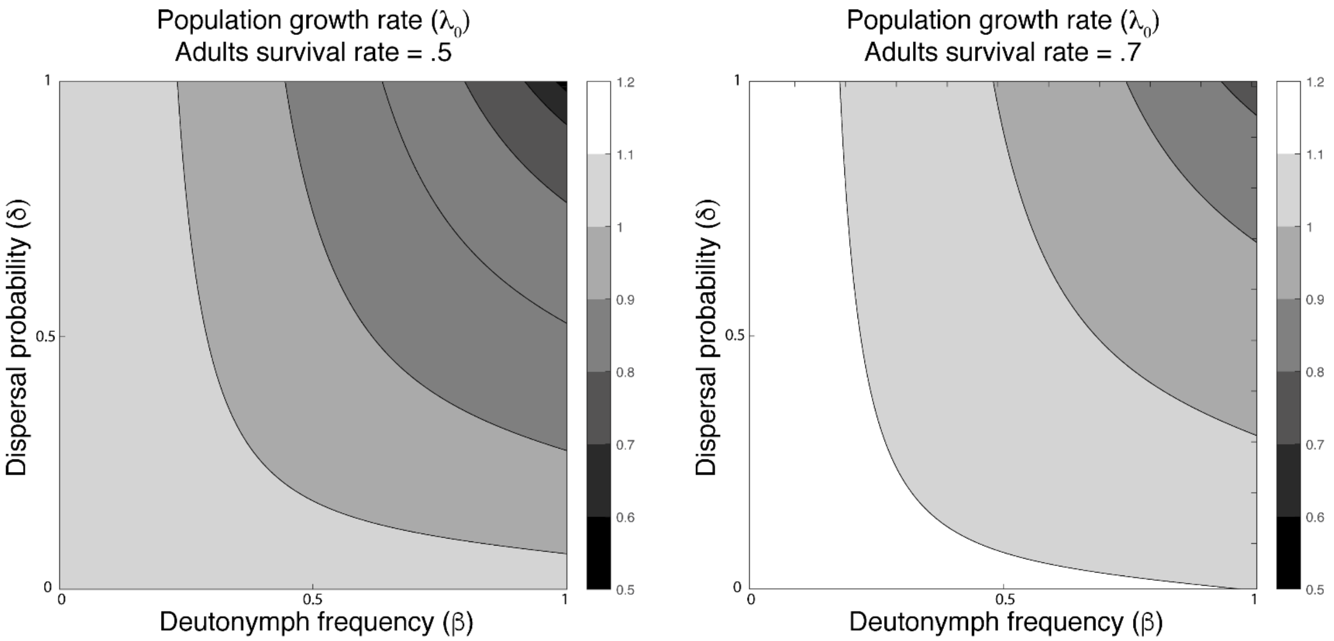


**Fig. S1**. Joint effect of increasing deutonymph frequency (β) and increasing dispersal probability ($\delta$) on population growth rate ($\lambda_0$). Dispersal probability and deutonymph frequency increase from 0 to 1 at 0.01 increments. Side bar indicates $\lambda_0$, with $\lambda_0$ increasing from black to white. The two panels indicate the response of $\lambda_0$ when adult survival rate within the DPM is set at 0.5 (left panel) and 0.7 (right panel).

#### Literature cited in the Appendix


Caswell, H. 2001. Matrix population models: Construction, analysis, and interpretation. Sunderland: Sinauer Associates, Massachusetts.

Deere, J. A., T. Coulson, and I. M. Smallegange. 2015. Life History Consequences of the Facultative Expression of a Dispersal Life Stage in the Phoretic Bulb Mite (*Rhizoglyphus robini*). PLoS ONE 10:e0136872.

Easterling, M.R., S.P. Ellner and P.M. Dixon. 2000. Size-specific sensitivity: Applying a new structured population model. Ecology 81: 694-708.

Ellner, S.P. and M. Rees. 2006. Integral Projection Models for Species with Complex Demography. The American Naturalist 167: 410-428.

R Core Team (2013). R: A language and environment for statistical computing. R Foundation for Statistical Computing, Vienna, Austria. URL http://www.R-project.org/.

Smallegange, I.M., J.A. Deere and T. Coulson. 2014. Correlative changes in life-history variables in response to environmental change in a model organism. American Naturalist 183: 784-797.
